# Streamlined, recombinase-free genome editing with CRISPR-Cas9 in *Lactobacillus plantarum* reveals barriers to efficient editing

**DOI:** 10.1101/352039

**Authors:** Ryan T. Leenay, Justin M. Vento, Malay Shah, Maria Elena Martino, François Leulier, Chase L. Beisel

## Abstract

Lactic-acid bacteria such as *Lactobacillus plantarum* are commonly used for fermenting foods and as probiotics, where increasingly sophisticated genome-editing tools are currently being employed to elucidate and enhance these microbes’ beneficial properties. The most advanced tools to-date require heterologous single-stranded DNA recombinases to integrate short oligonucleotides followed by using CRISPR-Cas9 to eliminate cells harboring unedited sequences. Here, we show that encoding the recombineering template on a replicating plasmid allowed efficient genome editing with CRISPR-Cas9 in multiple *L. plantarum* strains without a recombinase. This strategy accelerated the genome-editing pipeline and could efficiently introduce a stop codon in *ribB*, silent mutations in *ackA*, and a complete deletion of *lacM*. In contrast, oligo-mediated recombineering with CRISPR-Cas9 proved far less efficient in at least one instance. We also observed unexpected outcomes of our recombinase-free method, including an ~1.3-kb genomic deletion when targeting *ribB* in one strain, and reversion of a point mutation in the recombineering template in another strain. Our method therefore can streamline targeted genome editing in different strains of *L. plantarum*, although the best means of achieving efficient editing may vary based on the selected sequence modification, gene, and strain.

## INTRODUCTION

*Lactobacillus* represents a diverse and biotechnologically important genus of bacteria. As they naturally produce lactic acid, many *Lactobacilli* strains are commonly found in yogurts and other food products (Kailasapathy and Chin, 2000; Wang et al., 2004). This widespread usage has also pushed their development as ingestible probiotics to improve gut health. *In vivo* studies have demonstrated that members of the *Lactobacillus plantarum* can combat gut-residing infections in humans (Wullt et al., 2007). Recent work has also highlighted their ability to promote host growth under nutrient-limiting conditions in fruit flies and in mice (Schwarzer et al., 2016; Storelli et al., 2011). A significant part of these successful applications can be attributed to ever-advancing genetic tools. These tools have been used to elucidate how Lactobacilli genetics contribute to their desirable properties (Martino et al., 2018; Matos et al., 2017) and to forward-engineer strains as enhanced probiotics or to produce metabolites in different food products (Bron et al., 2007; Okano et al., 2018).

To date, a number of studies have established increasingly sophisticated genetic tools for *Lactobacilli* (Alegre et al., 2004; Aukrust, 1988; Bryan et al., 2000; Kleerebezem et al., 1997; Kullen and Klaenhammer, 2000; Thompson and Collins, 1996) that have created a framework for genome editing in this genus (Bron et al., 2007; Maguin et al., 1996, 1992; Okano et al., 2018, 2009). Recently, a scarless genome-editing system was developed in several Lactobacilli that relies on expressing a phage-derived single-stranded DNA recombinase to integrate oligonucleotides into the genome (van Pijkeren and Britton, 2012). This approach was further enhanced in the model bacterium *L. reuteri* by DNA cleavage with CRISPR-Cas9 (Oh and van Pijkeren, 2014). CRISPR-Cas9 is a two-component system comprising a guide RNA (gRNA) and the RNA-guided Cas9 nuclease from *Streptococcus pyogenes* (SpCas9) (Gasiunas et al., 2012; Jinek et al., 2012). The gRNAs are designed to be complementary to target DNA sequences flanked by a 3’ NGG protospacer-adjacent motif (PAM) based on a guide-centric orientation (Jinek et al., 2012; Leenay and Beisel, 2017), leading SpCas9 to cleave the DNA 3 bps upstream of the PAM. The gRNA can be an engineered single-guide RNA (sgRNA) or processed from a transcribed CRISPR array with the help of RNase III and a trans-acting CRISPR RNA (tracrRNA) (Deltcheva et al., 2011; Jinek et al., 2012). In bacteria, the traditional paradigm for oligo-based editing with CRISPR-Cas9 is that the oligonucleotide mutates the genomic site targeted by the sgRNA, and Cas9 only targets cells that did not undergo recombination. Due to the cytotoxicity of cleaving bacterial genomes with a CRISPR nuclease (Gomaa et al., 2014; Jiang et al., 2013; Leenay and Beisel, 2017; Vercoe et al., 2013), Cas9 effectively serves as a negative selection against unedited cells, thereby greatly boosting the frequency of successful editing.

While promising, these genome-editing tools have rarely been applied outside of model strains of Lactobacilli. One potential bottleneck relates to the recombinase. It normally must be under inducible control to limit unintended recombination, and the associated sensory proteins often must be co-expressed (e.g. NisR and NisK for nisin). Furthermore, the co-transformation of large quantities of the oligos and the CRISPR-Cas9 plasmid can reduce the overall transformation efficiency, thus not all cells will receive both the oligos and the plasmid. Recent work has begun to suggest that the recombinase can be dispensable by taking a different approach to recombineering. For instance, prior work in *E. coli* showed that RecA and the endogenous homologous recombination machinery could drive efficient recombineering between a plasmid harboring a double-stranded recombineering template and the genomic site targeted by SpCas9 (Cui and Bikard, 2016). Similarly, accumulating examples of Cas9-based genome editing in bacteria rely on a plasmid-encoded recombineering template without the use of a heterologous recombinase (Altenbuchner, 2016; Huang et al., 2015; Jiang et al., 2013). In a recent example in the model bacterium *Lactobacillus casei*, a single construct encoding a nicking mutant of SpCas9, a targeting CRISPR array, and a recombineering template achieved editing efficiencies up to 62% (Song et al., 2017). These examples suggested that Cas9-based editing could be achieved in *Lactobacillus plantarum* without the need for a recombinase.

Here, we developed a recombinase-free method of genome editing with SpCas9 for *L. plantarum*. The method relies on a recombineering template encoded on a replicated plasmid to achieve editing. We show that our method efficiently generated a premature stop codon into the riboflavin biosynthetic gene *ribB*, whereas oligo-mediated recombineering fails to generate this same edit. Then, we expand the design of the recombineering template to insert silent mutations in the acetate kinase gene *ackA* and a complete deletion of the β-galactosidase subunit gene *lacM*. Finally, we observed two instances where this editing approach failed--specifically through the recombineering template reverting to the WT sequence in the model *L. plantarum* strain WCFS1 and the generation of a consistent deletion within the target gene in the non-model *L. plantarum* strain NIZO2877. Our method therefore offers a streamlined means of genome editing in *L. plantarum*, although the availability of multiple editing methods are still necessary.

## RESULTS

### Constructs for recombinase-free genome editing use two*E. coli-Lactobacillus* shuttle vectors

We sought to simplify the editing pipeline in *L. plantarum* using a recombinase-free system (**Figure 1A**). To facilitate this pipeline, we generated two *E. coli-Lactobacillus* shuttle vectors to enhance the rate of cloning the final constructs (**Figure S1**). The CRISPR-Ca9 targeting plasmid encodes SpCas9, its tracrRNA, and a CRISPR array under the control of the P_*pgm*_ constitutive promoter (Duong et al., 2011). A CRISPR array was chosen over the standard single-guide RNA (Jinek et al., 2012) to avoid the need for a defined transcriptional start site and to allow multi-spacer arrays for multiplexed editing. New spacers are added by digesting unique cutsites within the processed region of an existing spacer (**Figure 1B**). This construct exhibited potent activity based on a large drop in the transformation efficiency when targeting a conserved site in the *rpoB* gene in both strains (**Figure S2**). The base recombineering-template plasmid was generated by adding *E. coli* replication components into a compatible shuttle vector (van Pijkeren and Britton, 2012) as well as a multi-cloning site for insertion of the recombineering template. Each plasmid was isolated from the methylation-free *E. coli* strain EC135 to boost the transformation efficiency (Zhang et al., 2012). In some cases, the plasmid was then passaged through the highly-transformable strain WCFS1 prior to transformation into other *L. plantarum* strains.

**Figure 1:**
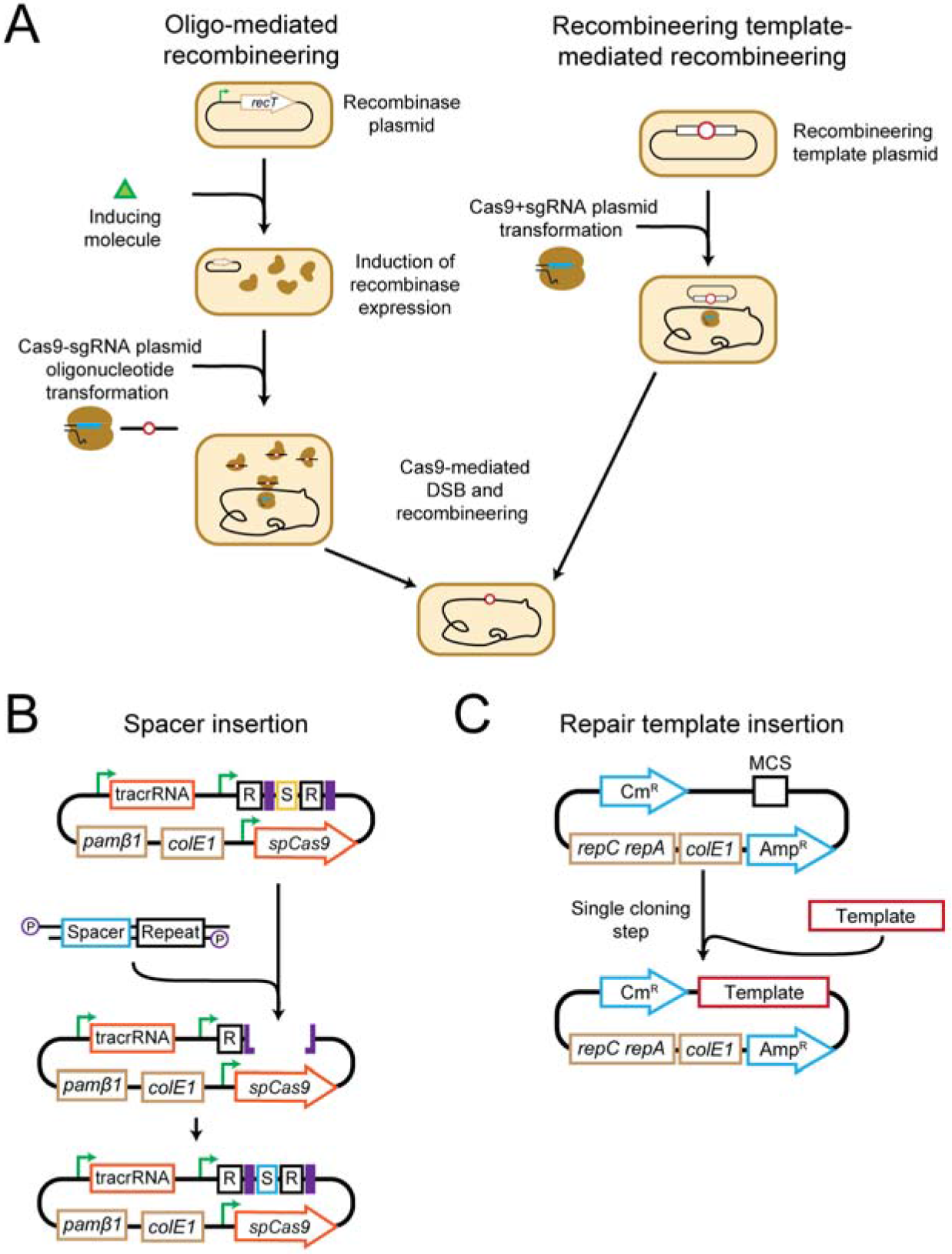
A pipeline for recombinase-free genome editing with CRISPR-Cas9 in *L. plantarum*. (**A**) Comparison of oligo-mediated recombineering and recombinase-free genome editing methods. (**B**) Cloning scheme to insert a new targeting spacer into the repeat-spacer-repeat array of the CRISPR-Cas9 construct. The base SpCas9 targeting plasmid is first digested with NotI and PvuI enzymes. A new spacer-repeat is designed as two oligonucleotides that are phosphorylated, annealed, and ligated into the digested backbone. All cloning is performed in *E. coli*. (**C**) Cloning scheme to insert a recombineering template into a plasmid that replicates in *E. coli* and *L. plantarum*. Gibson assembly is performed to ligate the amplified region of the targeted gene into the multiple cloning site of the base plasmid. Site-directed mutagenesis can be used to incorporate the desired change into the recombineering template. All cloning is performed in *E. coli*.

### The recombinase-free method efficiently generated a premature stop codon in *ribB* in *L. plantarum* WJL

As a test case for validating our recombinase-free genome editing constructs, we chose the moderately transformable *L. plantarum* strain WJL. This strain has been shown to provide a fitness benefit when added to germ-free fruit fly larvae or pre-weaned mice under nutrient-limiting conditions, and it could offer a chassis for engineered probiotics (Schwarzer et al., 2016; Storelli et al., 2011). We decided to introduce a premature stop codon into the *ribB* gene. This gene is crucial for the production of Vitamin B_2_, an essential nutrient that humans must obtain from their diet or their gut microbiota. Certain strains of *L. plantarum*, including the model strain WCFS1, do not contain a complete *ribB* operon (Kleerebezem et al., 2003) and require riboflavin for growth (Burgess et al., 2006). In contrast, the WJL strain contains a complete *ribB* operon (Martino et al., 2015a). We designed a novel spacer to target a site near the 5’ end of the *ribB* riboflavin synthase gene, and we designed a recombineering template with 1-kb homology on both sides of a premature stop codon that would incorporate into the 5’ end of the *ribB* gene. The stop codon was placed within the targeted PAM, thereby preventing any subsequent cleavage by SpCas9 once recombination occured (**Figure 2A**).

**Figure 2:**
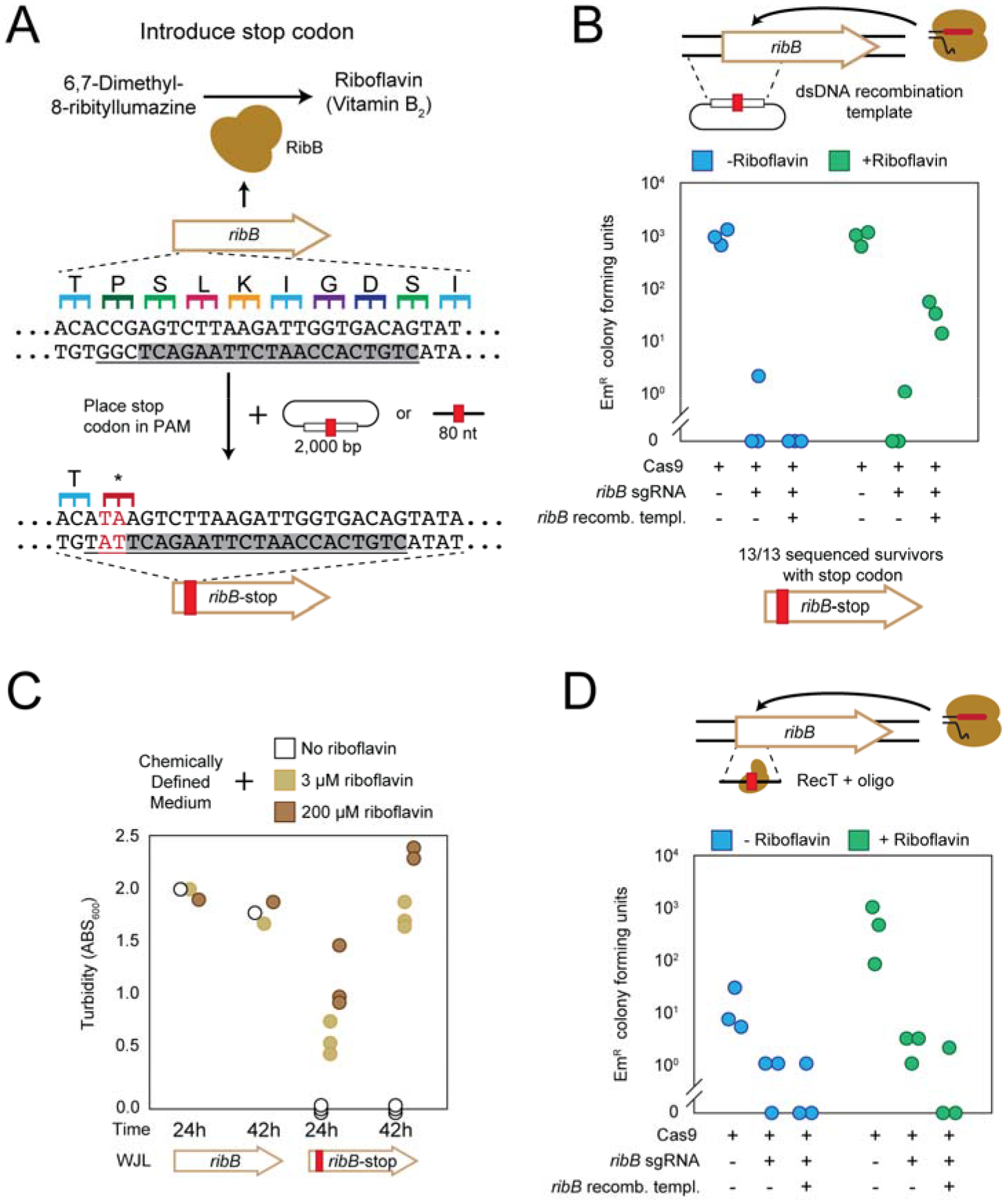
Enhanced editing of *ribB* by recombinase-free editing than by oligo-mediated editing in *L. plantarum* WJL. (**A**) A double-stranded repair template was designed along with a Cas9 targeting construct to generate an early stop codon in the *ribB* gene of *L. plantarum* WJL. Transformed cells were plated on Erythromycin MRS agar with or without supplementation of 200 µM riboflavin. (**B**) Surviving colonies were subject to Sanger sequencing for validation of the premature stop codon. Three confirmed mutants were grown on chemically defined medium (CDM) with the specified concentration of supplemented riboflavin to asses the phenotype of the riboflavin synthase knockout. (**C**) A single-stranded oligo was transformed along with the Cas9 targeting construct into *L. plantarum* WJL containing RecT recombinase induced with nisin. Cells transformed with the indicated plasmids were plated on Erythromycin MRS agar with/without supplementation of 200 µM riboflavin. Replicates represent independent experiments starting from separately validated mutant strains.

Editing was performed by first transforming the cells with the recombineering-template plasmid followed by the *ribB*-targeting plasmid. Cells were then plated on MRS with or without 200-µM riboflavin to test whether the mutants required riboflavin for growth (**Figure 2B**). We did not obtain any colonies in the absence of riboflavin, while in the presence of riboflavin we obtained ~28-fold fewer colonies compared to a no-guide control. Survivors were subjected to colony PCR using primers that bind the genome outside the region used for the recombineering template, thereby preventing any false-positives. Of the 13 sequenced survivors, all contained the intended premature stop codon. After clearing the two plasmids, we found that the survivors surprisingly grew in regular MRS broth, likely due to trace amounts of riboflavin in the undefined media. A chemically defined medium (CDM) was therefore used to assess the *ribB*-deletion phenotype (Hébert et al., 2004). Strains containing the premature stop codon exhibited negligible growth in the CDM medium without riboflavin and modest growth in CDM with trace amounts of riboflavin (**Figure 2C**). This phenotypic result is in line with the essentiality of riboflavin production via the *ribB* operon for cell growth in *L. plantarum* WJL. Overall, we found that efficient genome editing without a recombinase is possible in *L. plantarum* WJL.

### The oligo-based method failed to generate the premature stop codon in *ribB* in *L. plantarum* WJL

After successful demonstration of genome editing, we used the *ribB* site to evaluate how our our method compared to the previously established method of oligo-based editing with CRISPR-Cas9 (**Figure 1A**). We first generated a construct containing NisR, NisK, and the P_nisin_ for nisin-inducible expression of RecT (Bryan et al., 2000) (**Figure S1**). The recombineering activity of RecT was confirmed by transforming the *L. plantarum* strains WCFS1 and NIZO2877 with an oligonucleotide conferring rifampicin resistance following prior work (**Figure S2B**) (Garibyan et al., 2003; van Pijkeren and Britton, 2012). We then confirmed that oligo-mediated recombineering with CRISPR-Cas9 could select for the same edit in WJL and WCSF1, although WJL showed low editing efficiency (2/23 sequenced survivors contained the intended edit) (**Figure S2C,D**). Next, an 80-nt single-stranded oligonucleotide was designed to incorporate the same premature stop codon used with recombinase-free editing. We then transferred the RecT construct into WJL, induced with Nisin, and co-transformed the oligo and the *ribB*-targeting construct following the previously reported method (Oh and van Pijkeren, 2014). The co-transformation yielded only two erythromycin-resistant colonies on riboflavin-supplemented plates compared to one survivor on plates without additional riboflavin (**Figure 2C**), and these survivors contained the WT sequence. Thus, in this initial example, the recombinase-free method efficiently generated the desired edit while the oligo-based method failed to generate any edits.

### The recombinase-free method simultaneously introduced multiple point mutations in the *ackA* gene in WJL

We subsequently evaluated whether this method could generate other mutations within different genes of WJL. The acetate kinase gene *ackA* was chosen as its loss of function in *L. plantarum* NIZO2877 has been correlated to a fitness benefit in *Drosophila* models similar to the host fitness benefits mediated by *L. plantarum* WJL (Martino et al., 2018). We designed a spacer to target the *ackA* gene and a recombineering template that generates three unique silent mutations within the the seed region of the target (**Figure 3A**). Transformation of the *ackA*-targeting construct in a strain harboring the recombineering-template construct yielded an ~400-fold drop compared to the no-guide control (**Figure 3B**). This drop was noticeably larger than the ~28-fold drop observed when targeting *ribB* and mutating the PAM sequence (**Figure 2B**), suggesting some differential targeting activity when mutating the guide sequence versus the PAM. Following multiple editing attempts, we screened 20 colonies for the intended edit, where 19 contained all three silent mutations, while the remaining colony harbored the WT sequence. The generation of three point mutations in the WJL *ackA* gene demonstrated that the recombinase-free method can be expanded to incorporate multiple mutations in the seed region of the spacer.

**Figure 3:**
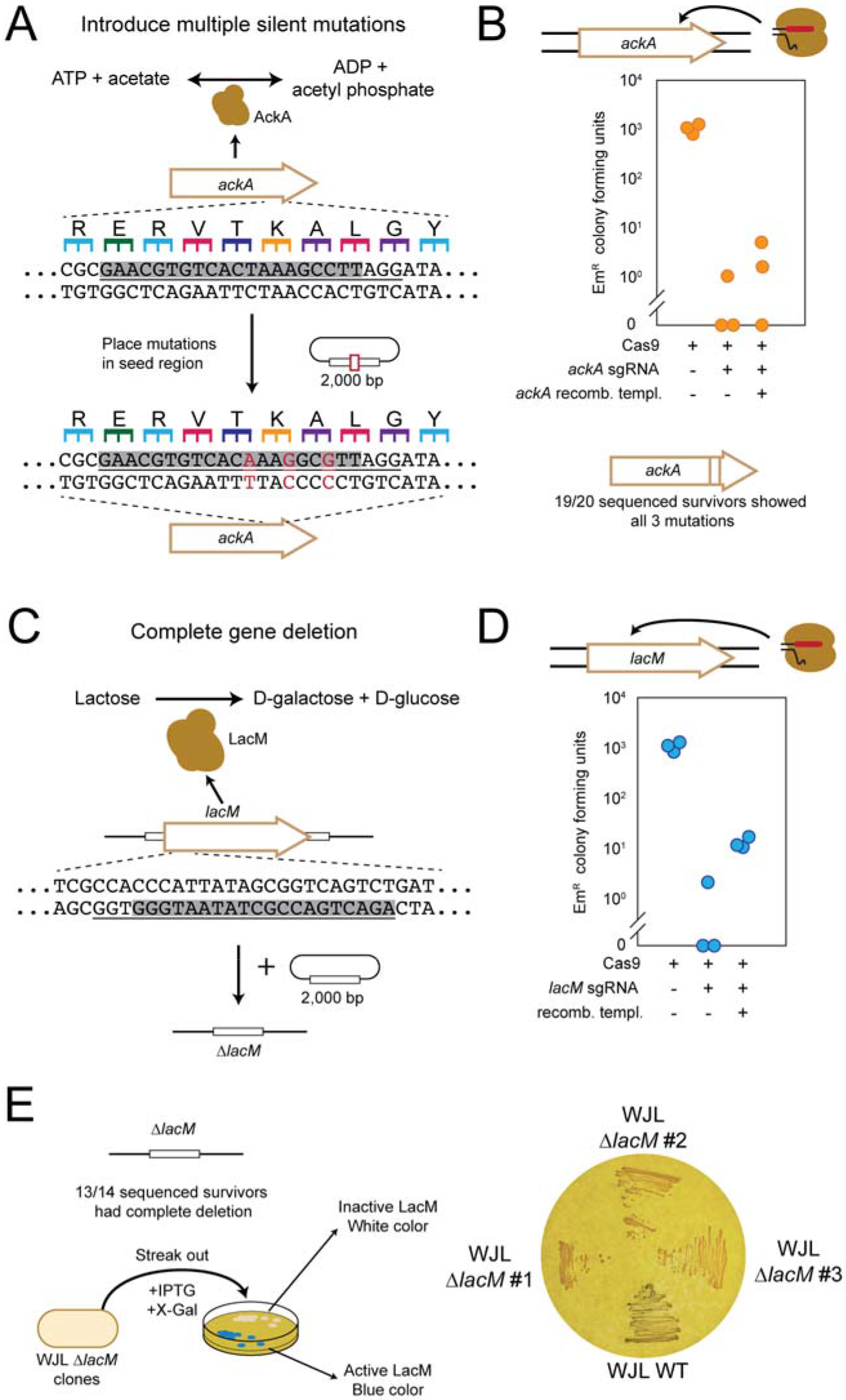
Recombinase-free genome editing can generate different types of edits in *L. plantarum* WJL. **(A)** The recombineering template was designed to incorporate three silent mutations into the acetate kinase *ackA* gene of WJL. The mutations fall within the seed region of the gRNA target site. (**B**) Cells transformed with the indicated plasmids were plated on erythromycin MRS agar, and survivors were subjected to Sanger sequencing to confirm the edited sequence. Replicates represent independent experiments starting from separate colonies. **(C)** The recombineering template was designed with 1-kb homology arms upstream and downstream from the start and stop codons, respectively, to generate a complete deletion of the *lacM* open-reading frame. The gRNA target site was within the deleted region. **(D)** Cells transformed with the indicated plasmids were plated on Erythromycin MRS agar. Replicates represent independent experiments starting from separate colonies. **(E)** Survivors were screened by colony PCR and gel electrophoresis and Sanger sequencing to to validate the 960-bp deletion. Three *lacM*-deletion mutants along with wild-type WJL were streaked on MRS agar containing Isopropyl β-D-1-thiogalactopyranoside (IPTG) and X-gal to check for a change in the colony color.

### The recombinase-free method deleted the entire *lacM* open-reading frame in WJL

We then explored whether our recombineering template-based method could yield a complete gene deletion. We chose the lactose mutase *lacM*, a 960-bp gene that expresses one of the subunits of β-galactosidase in *L. plantarum (Iqbal et al., 2010)*. The enzyme complex is responsible for breaking down lactose into D-galactose and D-glucose as well as producing galacto-oligosaccharides commonly used as prebiotics. For this reason, β-galactosidase activity is used as a marker to screen Lactobacilli with significant probiotic potential (Mandal and Bagchi, 2018). Importantly, the other subunit encoded by *lacL* cannot produce a fully functional beta-galactosidase enzyme if *lacM* is absent (Nguyen et al., 2007). Therefore, the recombineering template was designed to include 1 kb of homology upstream and downstream of the *lacM* open-reading frame, where recombination would result in a complete deletion from the start codon through the stop codon (**Figure 3C**). Transformation of the *lacM*-targeting construct into the recombination template-containing strain yielded an ~35-fold drop in the number of colonies compared to the no-guide control (**Figure 3D**), comparable to the CFU drop observed when mutating the PAM (**Figure 2B**). Of these 14 screened survivors, 13 harbored the intended *lacM* deletion. To phenotypically confirm successful deletion, we streaked mutants on MRS plates containing Isopropyl β-D-1-thiogalactopyranoside (IPTG), the synthetic inducer of the lac operon in *E. coli*, and X-gal, a compound that β-galactosidase hydrolyzes to generate a blue dye. Thus, cells containing a functional β-galactosidase should be blue, and cells expressing non-functional enzymes should be white. On the IPTG+X-gal plates, the three tested *lacM-*deletion mutants yielded white streaks while the WT strain yielded blue-ish streaks, confirming disruption of *lacM* (**Figure 3E**). The successful editing of *lacM* marks the third distinct gene that was successfully and efficiently edited using our recombinase-free method.

### The recombinase-free method revealed two potential failure modes

To investigate how well this recombinase-free genome editing method works in other *L. plantarum* strains, we attempted to generate the same stop codon in *ribB* in the less tractable strain NIZO2877 (**Figure S2A**) that generated ~10 fold fewer colonies compared to WJL after transformation with 5 µg of no-guide control. Once the recombineering-template plasmid was introduced into NIZO2877, we transformed the previous *ribB*-targeting plasmid and plated cells on MRS with or without supplemented riboflavin. Surprisingly, transformation of the *ribB*-targeting construct into the recombination template-containing strain yielded only ~1.8-fold fewer colonies compared to the no-guide control, and there were a comparable number of survivors on MRS plates with or without supplemented riboflavin (**Figure 4A**). As the original colony-PCR primers did not yield a band for any of the surviving colonies, we used a primers pair with a larger intervening distance. These wider flanking primers yielded a consistent ~1.3-kb deletion spanning upstream and into the 5’ end of *ribB* (**Figure 4A**) and partially overlapping with the recombineering template, including the Cas9 target site. The deletion was not necessarily due to failed editing activity in NIZO2877, as we recently and successfully applied the recombinase-free method in this strain to delete a codon within the *ackA* gene (Martino et al., 2018). That aside, the unintended deletion represents one means by which the Cas9-based editing can fail.

**Figure 4:**
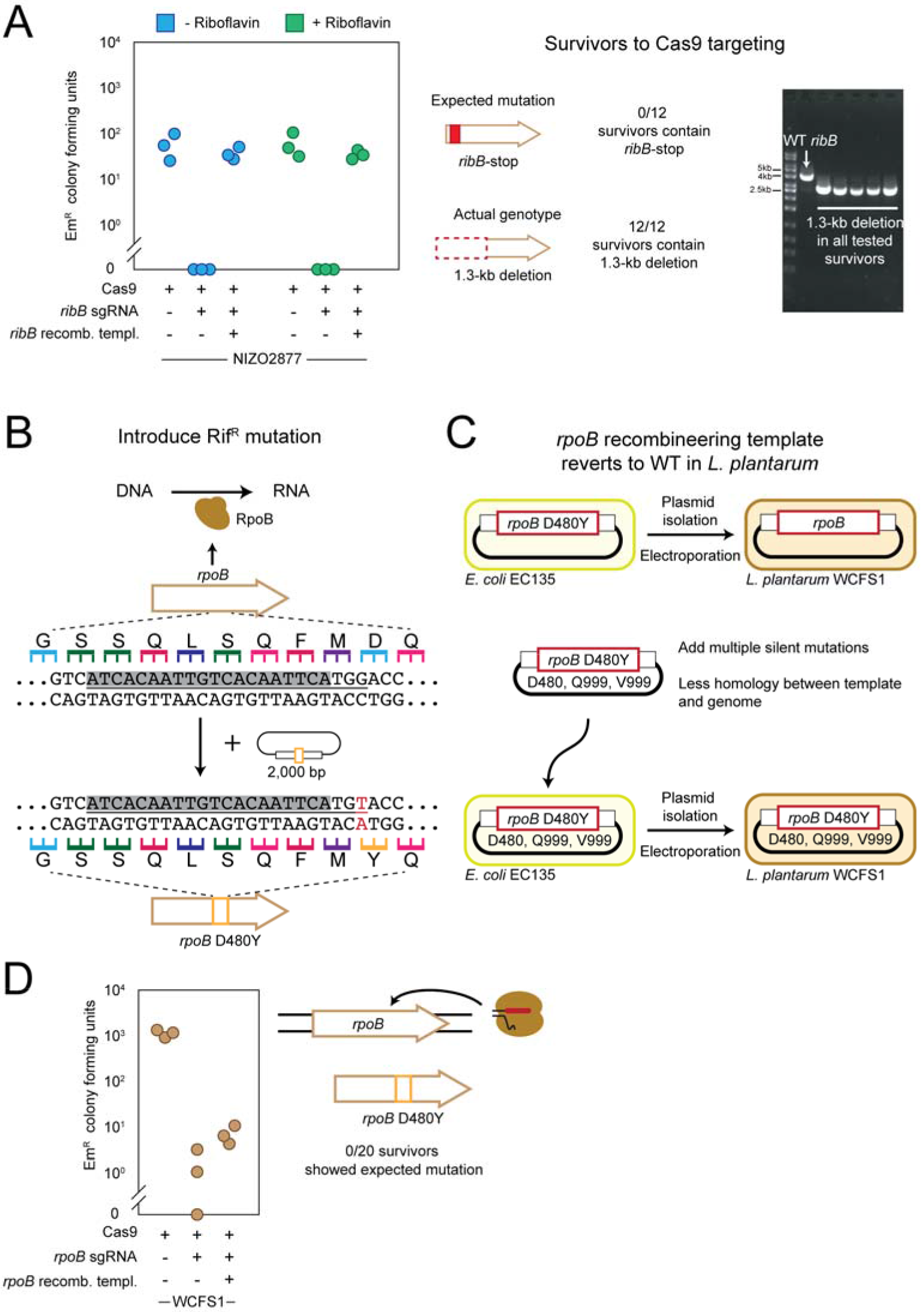
Instances of failed recombinase-free editing in *L. plantarum*. **(A)** The recombineering template and the *ribB*-targeting construct from Figure 2A was used to generate the premature stop codon in *ribB* in *L. plantarum* NIZO2877. Cells transformed with the indicated plasmids were plated on erythromycin MRS agar with or without supplemented 200-μM riboflavin. Lane 2 of the gel from colony PCR is from wild-type NIZO2877, and lanes 3-7 represent\ different survivor colonies. Replicates represent independent experiments starting from separate colonies. Sanger sequencing confirmed the cPCR results. **(B)** A recombineering template was designed to insert a point mutation conferring resistance to rifampicin into the *rpoB* gene of WCFS1. The mutation also disrupts the PAM in the gRNA target site. **(C)** The recombineering template reverted to the WT sequence when transformed into *L. plantarum* WCFS1 (right). Three silent mutations were subsequently cloned alongside the desired point mutation to prevent reversion of the recombineering template upon electroporation into WCFS1. **(D)** Cells transformed with the indicated plasmids were plated on erythromycin MRS agar. Survivors were subjected to Sanger sequencing to confirm the desired mutated sequence. Replicates represent independent experiments starting from separate colonies.

The recombinase-free method was also used to target the RNA polymerase *rpoB* gene in the model strain WCFS1, the earlier described test case to demonstrate editing with the oligo-based method (**Figure S2D**). The recombineering template was thus designed to include the same single point mutation conferring rifampicin resistance and flanked by 1-kb homology arms (**Figure 4B**). After transforming the previously used *rpoB*-targeting construct into cells containing this recombineering template, we screened 20 survivors and all contained the WT sequence. Further investigation revealed that the single point mutation within the recombineering template had reverted to the wild-type sequence after it was transferred from *E. coli* to WCFS1 (**Figure 4C**). Hypothesizing that recombination was occurring between the naturally occuring genomic sequence and our recombineering template, three additional silent point mutations were added around the rifampicin-conferring mutation to reduce the extent of homology between the template and the WCSF1 genome. Transferring the updated recombineering-template plasmid from EC135 into WCFS1 did not result in reversion of the point mutations, suggesting extensive homology may have been the cause. Subsequent transformation of the *rpoB*-targeting construct yielded surviving colonies, although screening 20 colonies only yielded the WT sequence (**Figure 4D**). These results revealed a unique failure mode of the recombinase-free editing method, where the template appeared to undergo recombination with the genome to lose the desired mutation. Even addressing this issue did not produce the desired edit, suggesting that other failure modes remain to be identified.

## DISCUSSION

We developed a method of Cas9-mediated genome editing in *L. plantarum* that relies on a double-stranded recombineering template rather than a heterologous recombinase and a single-stranded oligo. Our method efficiently produced various edits in three different genes, including introduction of a premature stop codon in *ribB*, generation of multiple silent mutations in *ackA*, and deletion of the entire *lacM* open-reading frame (**Figures 2, 3**). In one direct comparison with the oligo-based method editing *ribB* in WJL, we found that the recombinase-free method yielded efficient editing while oligo-mediated recombineering did not yield any edits. However, in another direct comparison editing *rpoB* in WCFS1, the oligo-method was successful whereas the recombinase-free method was not (**Figures 4B, S2**). Therefore, both methods can be utilized, although their efficiency may depend on the target and the strain.

As part of this work, we improved on the previously reported method for oligo-mediated recombineering with CRISPR-Cas9 (Oh and van Pijkeren, 2014). We placed the RecT, Cas9, the tracrRNA, and the CRISPR array into *E. coli-Lactobacillus* shuttle vectors to simplify and accelerate cloning. We also equipped the RecT plasmid with *nisR* and *nisK* so the plasmid could be used in strains lacking these genes. These constructs should aid others applying the oligo-based editing method.

Despite successful examples of recombinase-free editing, we did encounter two failed attempts at editing while applying this method. In one instance, a recombineering template was designed to generate a premature stop codon into the *ribB* gene of the non-model strain NIZO2877, but instead generated an ~1.3-kb genomic deletion (**Figure 4A**). Similar excisions have been observed when targeting genomic islands in other bacteria, where surviving cells circumvented to genome targeting by eliminating the targeted region (Selle et al., 2015; Vercoe et al., 2013). In a separate instance, we observed the reversion of a point mutation in the recombineering template to the wild-type sequence after passaging from *E. coli* to *L. plantarum*, and before introducing Cas9 (**Figure 4C**). It may be that introducing a point mutation of this essential gene on a shuttle vector somehow is cytotoxic, as growth of the recombineering template strain was slower compared to WT WCFS1 (data not shown). These failed instances of genome editing should be instructive as others perform genome editing with recombineering templates or explore ways of improving the method.

Looking ahead, there is ample room for improving CRISPR-mediated, recombinase-free genome editing in Lactobacilli. Although SpCas9 has been the standard for achieving genome editing in bacteria, the CRISPR nuclease Cas12a/Cpf1 is becoming more prevalent (Hong et al., 2018; Yan et al., 2017; Zetsche et al., 2015) and in once instance yielded efficient editing when Cas9 could not be introduced (Jiang et al., 2017). There are also other emerging approaches to enhance editing that could be incorporated to our method. For instance, the Cas9 and the gRNA can be placed under inducible control to ensure that a large population of cells possess CRISPR-Cas9 and the recombineering template prior to DNA induction (Reisch and Prather, 2015). The challenge is finely tuning Cas9 induction to ensure negligible leaky expression in the absence of inducer and sufficient expression to drive editing in the presence of inducer. Separately, using a nicking Cas9 in multiple bacteria including the model bacterium *Lactobacillus casei* was shown to increase editing efficiencies (Li et al., 2018; Song et al., 2017; Xu et al., 2015), presumably based on promoting homologous recombination without introducing a lethal double-stranded break. However, because of the lack of lethality, the technique often leaves unedited cells. These other options provide ways to further enhance editing in *L. plantarum* and other lactic-acid bacteria, even as they introduce additional engineering challenges.

Overall, our study simplified the machinery required for efficient genome editing in *L. plantarum* WJL, and provided the framework for achieving editing in other non-model strains of Lactobacilli (Martino et al., 2018). The insights we gathered while applying our method highlight that strain-dependent behavior is a major obstacle to streamlining genome editing in this diverse genus, as the best means of editing may vary based on the desired edit, gene, and target strain. Continuously-evolving CRISPR genome editing tools will likely drive enhanced editing capabilities in Lactobacilli, which should in turn expand our understanding of their genomic versaitility and open the door to improved and novel applications.

## MATERIALS AND METHODS

### Strains, plasmids, oligonucleotides

Table S1 contains descriptions and locations of every plasmid, strain, and oligonucleotide used for this study. *Lactobacillus plantarum* NIZO2877 and WJL were sent to us by Dr. Francois Leulier (Martino et al., 2015a, 2015b), and *L. plantarum* WCFS1 was sent to us from Dr. Nikhil U. Nair. Plasmids pJP042 and pJP005 were sent to us by the van-Pijkeren lab, and pMSP3545 (CN#46888) and pCas9 (CN#42876) were purchased from Addgene. Plasmids from this work will made available through Addgene.

### Plasmid generation

To generate pRecTNisRK, pJP005 was digested with XbaI and HindIII and the *nisR* and *nisK* genes were amplified with oRL9-10 and digested with XbaI and HindIII. The ligation was performed with 200 ng of digested backbone with a 3:1 molar excess of insert and was subsequently ethanol precipitated and transformed into *L. plantarum* WCFS1. Colonies were screened with oligos oRL7-8 by Miniprep (Zymo CN# D4036) of plasmids after cells were lysed with 20 ng/mL of Lysozyme for 30 minutes.

The CRISPR-Cas9 plasmids were created by taking the base pMSP3545 shuttle vector and pCas9 and amplifying each fragment with oRL1-oRL4. These PCR fragments were assembled by Gibson assembly kit (NEB CN# E2611S) (Gibson et al., 2009). This created the non-targeting p3545Cas9 control plasmid containing Cas9 and the tracrRNA. The targeting repeat spacer array was designed as a gBlock (3545_RSR_gBlock) and amplified with oRL5-6. The p3545Cas9 backbone was digested with XbaI and PstI and the fragments were assembled together with Gibson. This created the pCas9_RSR plasmid. Subsequent spacers were cloned in by digesting the backbone with PvuI and NotI, and annealing two oligonucleotides containing these same overhangs (oRL11-oRL12, oRL19-20, oRL27-28, oRL33-34) and ligating the fragments together.

The recombineering template shuttle plasmid (RLShut) was generated by amplifying the pJP005 backbone with oRL15-16, amplifying the ColE1 origin and *bla* resistance gene from pBAD18 with oRL13-14, then stitching the two pieces together by Gibson assembly. Functional clones were screened through a chemical transformation assay into *E. coli* DH5-alpha. Double-stranded DNA (dsDNA) recombineering templates were inserted into this shuttle vector by amplifying the desired repair sequence (oRL17-18, oRL21-22, oRL25-26, oRL31-32), digesting the PCR fragment and RLShut backbone with SpeI and SacI, and ligating the fragments together. If the PCR fragment was amplified from the wild-type genome, Q5 site directed mutagenesis (NEB CN# E0554S, oRL23-24) was utilized to create small nucleotide changes. After successful clones were generated in *E. coli* DH5-alpha, the plasmid was passaged through *E. coli* EC135, then *L. plantarum* WCSF1, and then transferred into the intractable *Lactobacillus* strain.

### Standard growth conditions

All *L. plantarum* strains were grown on MRS liquid broth (BD CN# 288130) and MRS agar (BD CN# 288210) and incubated at 37°C without shaking in 14-mL polypropylene tubes. Antibiotic concentrations in *L. plantarum* were as follows: rifampicin (25 µg/mL), chloramphenicol (10 µg/mL), and erythromycin (10 µg/mL). For the *ribB* knockout, MRS liquid broth was supplemented with and without 200 µM riboflavin. For *lacM* knockout, MRS agar was supplemented with 100 µM IPTG and 3 µL of X-gal per 1 mL of MRS.

All *E. coli* propagation was performed in LB medium (10 g/L NaCl, 5 g/L yeast extract, 10 g/L tryptone) while being shaken at 250 rpm at 37°C. Plasmids were maintained at the following antibiotic concentrations: erythromycin (50 µg/mL for liquid cultures, 300 µg/mL for plates), chloramphenicol (34 µg/mL), and ampicillin (50 µg/mL).

### Electroporation protocol for *L. plantarum*

Electroporation of *L. plantarum* was adapted from numerous protocols (Alegre et al., 2004; Aukrust, 1988; Spath et al., 2012; Thompson and Collins, 1996). A *L. plantarum* colony was picked from a plate after 48 hours of growth and placed into 3 mL liquid culture containing necessary antibiotics for 18 hours. 1 mL of this culture was back-diluted into 25 mL of MRS containing 0.41 M glycine and any necessary antibiotics. Outgrowth were performed in 50 mL falcon tubes (VWR CN#21008-212) to prevent aeration of the bacteria. These tubes were cultured at 37°C and 250 RPM until the OD_600_ was approximately 0.85 (~3.5 hours). If nisin was used to induce *recT* expression, 2 ng/mL of Nisin was added when the OD_600_ was approximately 0.7 (after ~3 hours of outgrowth), and cultures were shaken until they reached an OD_600_ of 0.85. Then, cells were centrifuged at 5,000 RPM for 10 minutes at 4°C to collect the pellet, and washed twice with 5 mL of 10 mM MgCl_2_. After transferring to a new 50 mL tube, cells were and washed with 5 mL of SacGly (10% glycerol with 0.5M sucrose) (Alegre et al., 2004; Spath et al., 2012). Finally, *L. plantarum* cells were washed in 1 mL of SacGly and centrifuged at 20,000 rpm for 1 minute. The supernatant was removed and the final pellet was resuspended in 500 µL of SacGly. For all transformations, 60 µL of this suspension was added to a 1-mm gap cuvette and transformed at 1.8 kV, 200 Ω resistance, and 25 µF capacitance. DNA concentrations used for transformations are reported in the main text. On average, 5 µg dsDNA and 10 µg of ssDNA oligo was transformed into *Lactobacillus* cells. Following electroporation, 1 mL of MRS broth was added to the cuvette and transferred to a sterile tube and incubated at 37°C without shaking for 4 hours. 250 µL of this recovery was then plated on MRS agar with necessary antibiotics. Any dilutions prior to plating was done in MRS media.

### Colony PCR

Colony PCR was performed by picking a single colony into 20 µL of 20 mM NaOH and incubating at 98°C for 20 minutes. These tubes were then microwaved for 1 minute with the cap open, and this mixture was diluted 1:10, and 5µL was added to the reaction. This was combined with NEB OneTaq HotStart 2x MM (CN# M0484).

### Plasmid clearance

In order to remove plasmids from mutated *L. plantarum* strains, cells were passed through multiple rounds of inoculation in MRS broth supplemented with appropriate nutrients and without any antibiotics. Once cells exhibited no antibiotic resistances in media and on agar plates, they were screened through colony PCR to ensure mutation remained intact.

### RibB phenotype assessment

Mutant and wild-type *L. plantarum* cells were grown on chemically defined media (Hébert et al., 2004) with and without 200-uM riboflavin to assess the effect of the *ribB* knockout. An OD_600_ measurement was taken after 24 and 42 hours on a spectrophotometer (Thermo Fisher CN# ND2000) to assess growth.

### LacM phenotype assessment

Strains of *L. plantarum* WJL with a clean *lacM* deletion were grown on MRS agar with 100-µM of Isopropyl β-D-1-thiogalactopyranoside (IPTG) and 3 µL of ready-to-use X-gal (Thermo CN# R0941) per 1 mL MRS. After 48 hours, colonies were checked for phenotypic color changes compared to wild-type WJL growth on the same plate.

## ACKNOWLEDGEMENTS

Plasmid pJP005 was a gift from the van Pijkeren lab. Plasmid pMSP3545 was a gift from Gary Dunny (Addgene plasmid # 46888) and plasmid pCas9 was a gift from the Luciano Marraffini (Addgene plasmid # 42876). This work was support by a CAREER award from the National Science Foundation (MCB-1452902 to C.L.B.), and a European Research Council starting grant (FP7/2007-2013-N°309704 to F.L), and the European Union’s Horizon 2020 research and innovation program under the Marie Sklodowska-Curie grant agreement (N°659510 to M.E.M.).

## AUTHOR CONTRIBUTIONS

R.T.L., J.M.V., and C.L.B. designed the experiments. R.T.L, J.M.V., M.S., and C.L.B. analyzed and formatted all data. M.E.M. and F.L. developed riboflavin for editing and phenotypic measurements. R.T.L. generated the figures and J.M.V., M.S., and C.L.B. wrote the manuscript. All authors read and approved the manuscript.

## Abbreviations

CRISPR: Clustered Regularly Interspaced Short Palindromic Repeats
Cas: CRISPR-associated
sgRNA: single-guide RNA
RT: recombineering template

